# Correlation analysis between drug resistance and virulence genes of carbapenem-resistant *Acinetobacter baumannii* In Western Guilin

**DOI:** 10.1101/2022.09.28.510023

**Authors:** Lijun Xiong, Xiaofeng Huang, Huoying Chen, Zhenyu Liu, Di Wang, Guifen Zeng, Shan Mo, Chuandong Wei

## Abstract

**Backgound:** In order to detect the resistance of *Acinetobacter baumannii* to β-lactam antibiotics in western Guilin, analyze the reasons for its resistance, and provide laboratory basis for clinical treatment; method for the β-lactamase gene of *Acinetobacter baumannii*, and to explore the relationship between the presence of the β-lactamase gene of *Acinetobacter baumannii* and multidrug resistance.

**Methods:** From November 2020 to June 2022, 78 non-repeated *Acinetobacter baumannii* strains from the Second Affiliated Hospital of Guilin Medical University were collected, and their resistance to carbapenem antibiotics was analyzed. PCR was used to detect resistance-related genes (*bla*_OXA-23_, *bla*_OXA-51_, *bla*_OXA-58_, *bla*_OXA-24_, *bla*_IMP_ and *bla*_SIM_) and virulence genes (*cusE, abaI* and *bfmS*). To analyze the relationship between drug resistance and virulence genes of *Acinetobacter baumannii*.

**Results:** Most of the 165 non-enterobacteriaceae bacteria we collected from the hospital were *Acinetobacter baumannii* and *Pseudomonas aeruginosa*, and most of the specimen types were from sputum and secretions, and most of them were from respiratory and critical care diseases area and intensive care unit, among them, there are 78 strains of *Acinetobacter baumannii* studied this time. After screening, 78 *Acinetobacter baumannii* strains were resistant to Cefazolin, Ampicillin, Amoxicillin/Clavulanate, Aztreonam, Chloramphenicol, Tetracycline, Cefotaxime, Ampicillin/Sulbactam, Piperacillin, Gentamicin, Ciprofloxacin, Levofloxacin, Moxifloxacin, Cefepime, Ceftazidime, Piperacillin/Tazobactam, Compound Sulfonamide, Meropenem, Imipenem, Amikacin, Polymyxin, resistance rates were 98.72%, 98.72%, 96.16%, 96.15%, 96.15%, 66.67%, 60.26%, 58.98%, 58.97%, 57.69%, 57.69%, 57.69%, 57.69%, 57.69%, 56.41%, 55.13%, 53.85%, 52.56%, 52.56%, 50%, 0%. Among them, 17 strains with drug resistance gene *bla*_OXA-51_ were detected,accounting for 21.79%; drug gene *bla*_OXA-23_, *bla*_OXA-24_, *bla*_IMP_, *bla*_OXA-58_, *bla*_SIM_ gene were not detected; 17 strains with virulence gene *bfmS* was detected, accounting for 21.79%; *abaI, csuE* virulence gene was not detected.

**Conclusion:** *Acinetobacter baumannii* in the hospital is highly resistant to carbapenems, mainly carrying *bla*_OXA-51_ resistance genes. Drug resistance is closely related to the virulence gene *bfmS*.

## Introduction

*Acinetobacter baumannii* is a major hospital-associated pathogen that causes a range of diseases, including respiratory, bloodstream, urinary tract, surgical site, and wound infections ^[1]^. This bacterium is an aerobic gram-negative coccus with relatively few known virulence factors compared to other gram-negative pathogens.Under the condition of weakened immunity, opportunistic infections in these parts of the human body are caused. More seriously, *Acinetobacter baumannii* is widely distributed in the hospital environment and can survive for a long time. The hospital-acquired infection of critically ill patients in the neonatal department can also be widely spread through the hands of medical staff, contaminated medical equipment and air, resulting in outbreaks of nosocomial infections.Because of its strong colonization ability and complex and variable drug resistance mechanism, *Acinetobacter baumannii* has brought difficulties to clinical anti-infective treatment and curbing nosocomial spread.This species also has a remarkable propensity for the rapid acquisition of resistance to an extensive range of antimicrobial agents. They can exhibit a major resistance profile, including carbapenems and other β-lactam antibiotics, leaving clinicians with limited therapeutic options.^[2]^

*Acinetobacter baumannii* has a tendency to develop resistance to a variety of antibacterial drugs, and the treatment of infections with highly resistant strains can be extremely difficult ^[3,4]^. Its ability to acquire various antimicrobial resistance genes has made it a very successful nosocomial pathogen, which has been associated with increased in-hospital mortality ^[5]^. *Acinetobacter baumannii* isolates are on the rise in Asian countries ^[6]^.

Previous studies have compared the distribution of genes associated with virulence and antimicrobial resistance to understand phenotypic differences between isolates ^[7–9]^. All available isolates from patients with *Acinetobacter baumannii* bacteremia were sequenced for genome-wide association studies to epidemiologically characterize genomic changes during the study period and to investigate the clinical significance of *Acinetobacter baumannii* genomic features. ^[10]^ These studies of molecular-level characterization, including antimicrobial resistance determinants and virulence factors of *Acinetobacter baumannii* isolates, can provide insights into how organisms evolve over time and may implicate Clinical epidemiological significance of somatic genomic features.

In recent years, the U.S. Centers for Disease Control and Prevention has designated multidrug-resistant Acinetobacter as an urgent threat pathogen, driving ongoing public health surveillance and preventive actions. In addition, the World Health Organization has included carbapenem-resistant *Acinetobacter baumannii* on the list of bacteria that pose the greatest threat to human health, and has prioritized research and development of new antimicrobial therapies.Therefore, the purpose of this study was to: (1) evaluate the clinical characteristics and outcomes of patients infected with carbapenicillin-resistant *Acinetobacter baumannii,* and (2) elucidate the mechanisms underlying carbapenicillin resistance in *Acinetobacter baumannii* strains and toxin analysis, and (3) to provide solutions for clinical drug use, so that the drug resistance of *Acinetobacter baumannii* can no longer affect nosocomial infections.

## 1 Materials and methods

### 1.1 Resurrection Strain

Sources of strains 78 non-repeated carbapenem-resistant *Acinetobacter baumannii* strains in our hospital from November 2020 to June 2022 were collected, that is, repeated isolates from patients with the same specimen were excluded. Use the Laboratory Information Management System to find the required strain information, associate the strain storage location according to the relevant strain information, and use this information to retrieve Non-Enterobacteriaceae bacteria from the −80 °C bacterial library. After sterilizing the tweezers under the lamp, clip out the paper and inoculate it on the MH medium, then use the inoculation loop to streak the three areas, and then place it in a 35°C incubator for 18-24 hours. The next day, the MH medium was taken out, and the presence or absence of miscellaneous bacteria was observed. The colonies with miscellaneous bacteria were purified and each strain was identified separately to obtain the target strain.

### 1.2 Strain Passage

#### 1.2.1

The target strains were 78 strains of *Acinetobacter baumannii* that might carry genes. First, the strains stored in the −80°C refrigerator by the paper method were passaged to the blood plate, and the three-zone line was made. This is the first generation. It was incubated in a 35°C incubator for 18-24 hours, and a single colony was picked the next day. The second-generation strains, the method and conditions are the same as the first-generation strains, and the second-generation strains obtained can be used for drug susceptibility testing.

#### 1.2.2 Preparation of bacterial suspension

use sterile physiological saline to prepare a bacterial suspension with a McFarland turbidity of 0.5-0.6, and add 1 ml of the bacterial suspension to 49 ml of sterile physiological saline with a sample gun. Mix well to obtain a bacterial suspension with a CFU of 3×10^6^.

#### 1.2.3 Loading

The mixed bacterial suspension was added to BD Phoenix™ M50, and an ATCC25922 quality control was also added. After incubating at 35°C for 18-24 hours, observe whether the quality control has passed. After the quality control has passed, the results of the drug susceptibility plates of other samples can be read.

### 1.3 Drug Susceptibility Test

The identification strips were used to identify, and the *Acinetobacter baumannii* was tested for drug susceptibility by the broth microdilution method, and the imipenem or meropenem-resistant strains were verified by the drug susceptibility disk diffusion method. Antibiotics used were Gentamicin, Cefazolin, Ceftazidime, Cefotaxime, Cefepime, Aztreonam, Ampicillin, Piperacillin, Amoxicillin/Clavulanate, Ampicillin/Sulbactam, Piperacillin Lacillin/Tazobactam, Compound Sulfa, Chloramphenicol, Ciprofloxacin, Levofloxacin, Moxifloxacin, Tetracycline, Amikacin, Imipenem, Meropenem, Polymyxa. The drug susceptibility results were determined according to the 2021 standard of the American Clinical and Laboratory Standards Institute (CLSI), and the drug susceptibility results of tigecycline were based on the U.S. Food and Drug Administration (Food and Drug Administration, FDA) recommended breakpoints.

Reference basis: use the BioMerieux company Active pharmaceutical ingredient(API) identification strip for determination, select the strains with imipenem or meropenem minimum inhibitory concentration(MIC)≥8μg/mL, and then use the KB(Disk diffusion method for drug susceptibility testing) method to screen (meropenem inhibition zone diameter ≤14mm or imipenem inhibition zone Diameter ≤ 19mm), 78 carbapenem-resistant *Acinetobacter baumannii* strains were finally screened out, cultured and stored for later use. The quality control strains are *Escherichia coli* strain (ATCC 25922), *Staphylococcus* aureus strain (ATCC 25923), *Klebsiella* pneumoniae strain (ATCC 700603) and *Pseudomonas* aeruginosa strain (ATCC 27853), all from the drug resistance monitoring of Guangxi Zhuang Autonomous Region center.

### 1.4 DNA extraction and PCR amplification

#### 1.4.1 DNA extraction

Take a 1.5ml EP tube, add 200-500ul of double-distilled water, take the target strain into a group of ten strains, pick equal amounts of colonies on each MH plate, and grind them into an EP tube to form uniform bacteria. Suspension (the bacterial concentration should not be too high), and mark the sample number on the tube. The EP tube containing the bacterial liquid was placed on the float and boiled in boiling water for 10 min, and the centrifugation speed was 13000 rpm, and the centrifugation was performed for 10 min. Take the supernatant as the template DNA source.

#### 1.4.2 PCR amplification

The polypeptide amplification system is as follows: PCR reaction contains 2 μL DNA, 2 μL forward primer, 2 μL reverse primer, 4 μL ddH_2_O and 10 μL Dream Taq Green PCR Master Mix(2×). Amplification conditions is shown in Table 1. The primers used are shown in Table 2. PCR products were subjected to agarose gel electrophoresis(2%), a constant voltage of 120V, and an electrophoresis time of 40min. The amplified positive products were sent to Bgi Genomics Co., Ltd(Shenzhen) sequencing, and the sequences were compared with those published in the NCBI database.

**Tab. 1.**
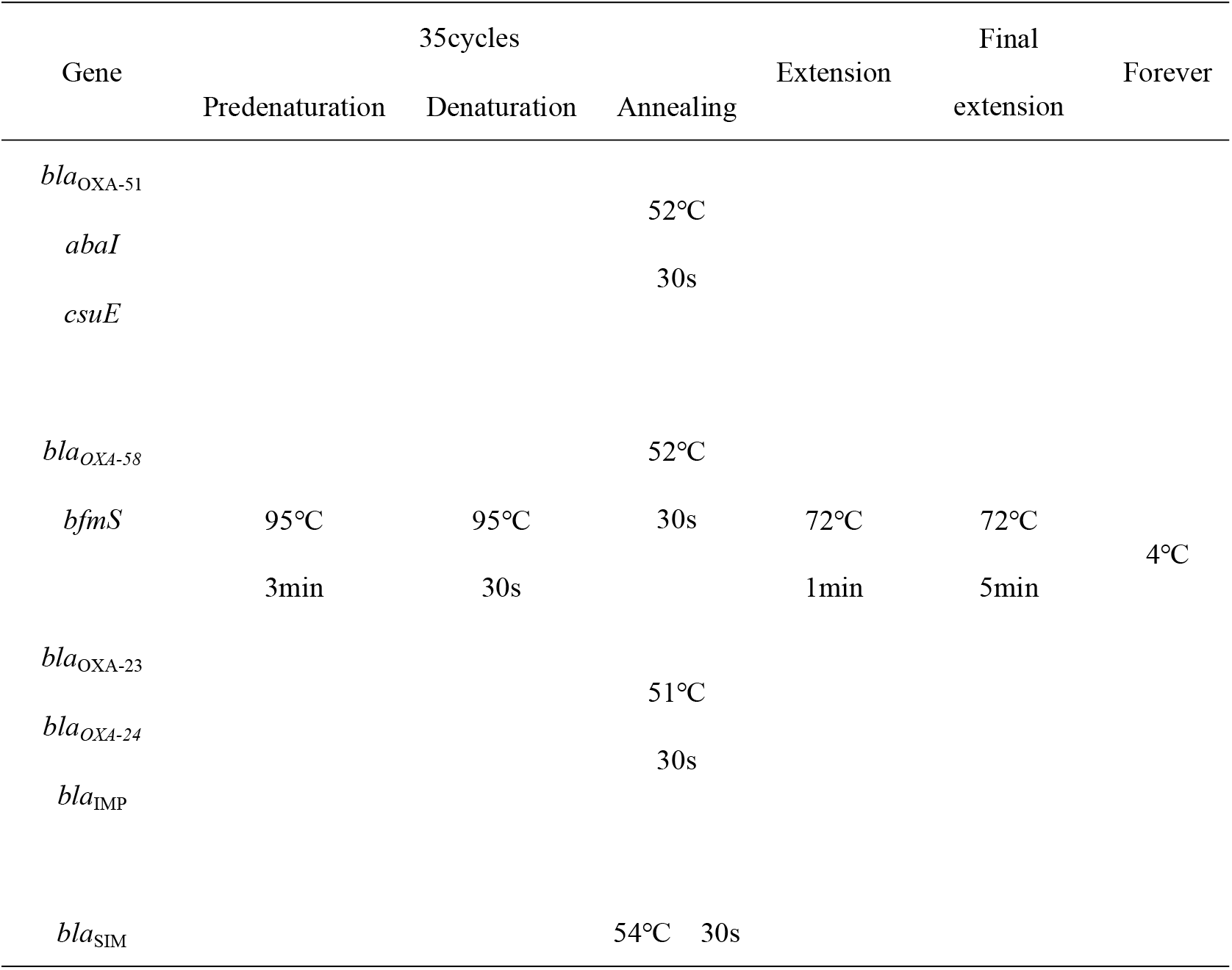
PCR amplification conditions

**Tab. 2.**
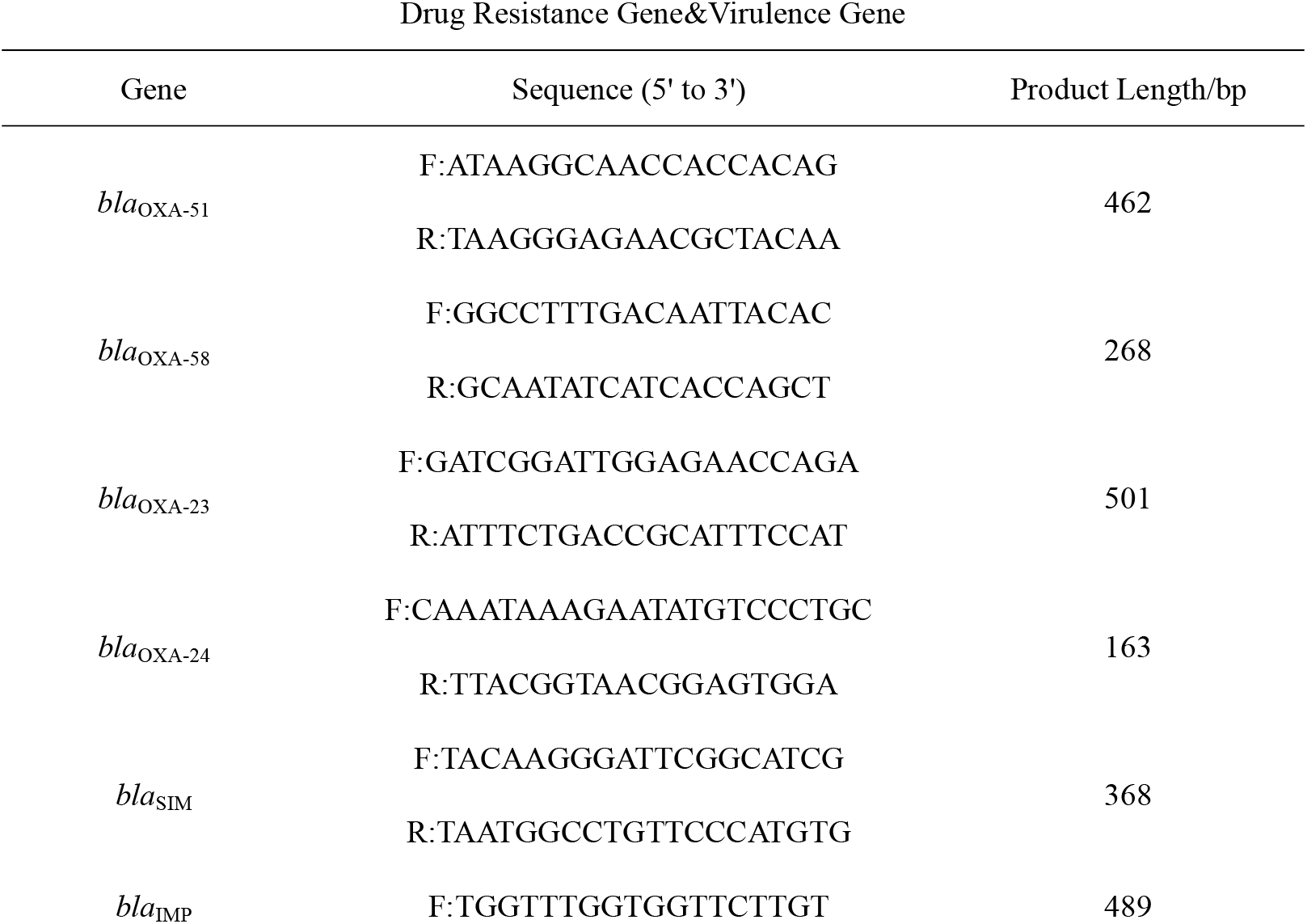
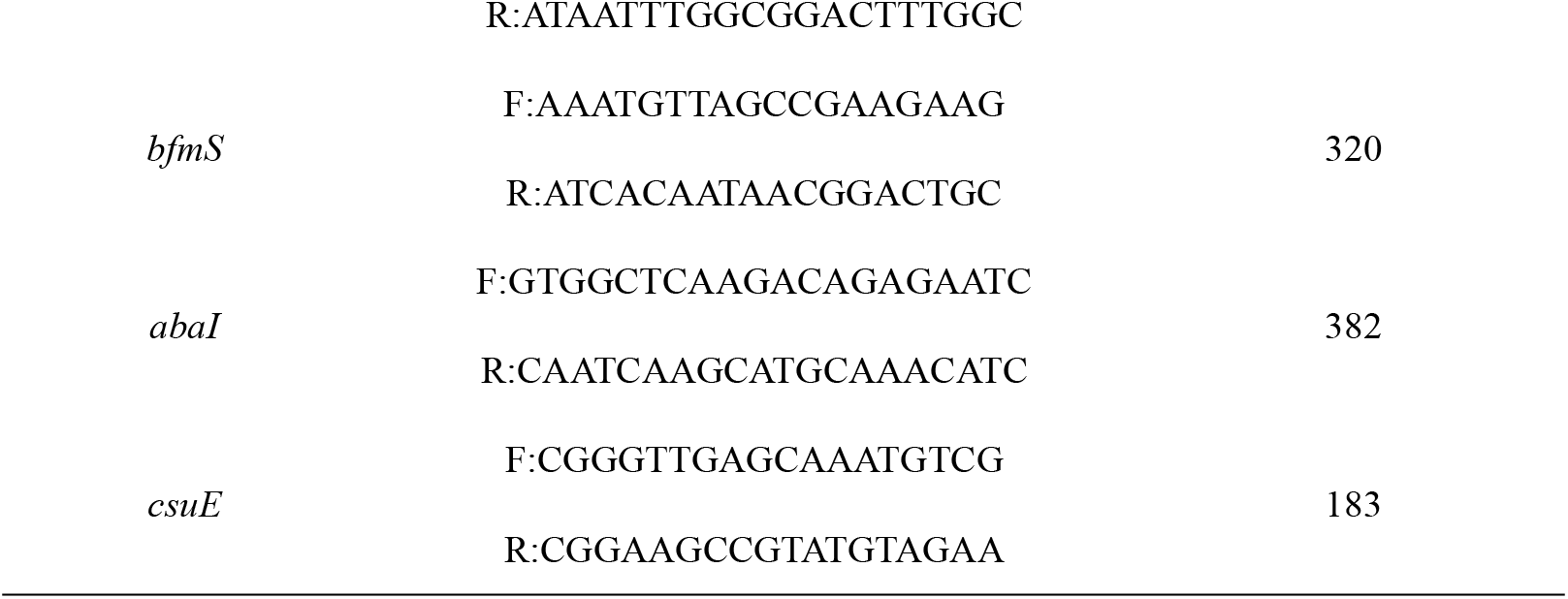
PCR primer sequence

### 1.5 Statistical analysis

SPSS 21.0 was used for data processing and analysis. The count data were expressed as percentage (%), and the *χ*^2^ test was used. P<0.05 indicated that the difference was statistically significant.

## 2 Result

### 2.1 Clinical features

The bacterial species composition (165 non-Enterobacteriaceae bacteria contained 18 different species, among which *Acinetobacter baumannii* accounted for 39.3%, Pseudomonas aeruginosa accounted for 35%, Stenotrophomonas maltophilia accounted for 5.7%, Mucoid Pseudomonas aeruginosa accounted for 5%, and the other species did not exceed 3%.)

Distribution of bacterial departments (140 non-Enterobacteriaceae bacteria were collected from 21 clinical departments, of which 28.6% came from the respiratory and critical care ward, 26.4% from the intensive care ward, and 6.4% from the dermatology ward, 6.4% came from the cerebrovascular disease ward, and the other collection points collected no more than 5%.)

Specimen type distribution (140 non-Enterobacteriaceae bacteria were derived from various clinical specimen types, 56.4% were from sputum specimens, 18.6% were from secretion specimens, 7.1% were from bronchoalveolar lavage fluid specimens, and the number of remaining specimen types are not more than 5%.), clinical characteristics of 78 patients with *Acinetobacter baumannii* see Table 3.

**Tab. 3.** Clinical characteristics of 78 patients with *Acinetobacter baumannii*

| Gender | Total number | Source | Total number | Department | Total number |
| --- | --- | --- | --- | --- | --- |
| Male | 52 | Sputum | 54(69.23%) | Intensive Medicine | 29(37.18%) |
| Female | 26 | Secretion | 8(10.26%) | Respiratory and critical care unit | 18(23.08%) |
| Age (years) | Total number | bronchoalveolar lavage fluid | 6(7.69%) | Cerebrovascular disease | 9(11.54%) |
| 0–20 | 3 | bile | 3(3.85%) | Neurosurgery | 5(6.41%) |
| 21–50 | 12 | Other | 7(8.97%) | Hepatobiliary and pancreatic surgery | 3(3.85%) |
| 51–70 | 30 |  |  | Orthopedic | 3(3.85%) |
| 70–more | 33 |  |  | Other | 11(14.10%) |

This study was approved by the Ethics Committee of the Second Affiliated Hospital of Guilin Medical College of Guangxi Zhuang Autonomous Region. Written informed consent was obtained from each patient or volunteer prior to sample collection. All samples were stored at −80°C until further experiments.

### 2.2 Drug resistance analysis of *Acinetobacter baumannii*

78 *Acinetobacter baumannii* strains were resistant to Cefazolin, Ampicillin, Amoxicillin/Clavulanate, Aztreonam, Chloramphenicol, Tetracycline, Cefotaxime, Ampicillin/Sulbactam, Piperacillin, Gentamicin, Ciprofloxacin, Levofloxacin, Moxifloxacin, Cefepime, Ceftazidime, Piperacillin/Tazobactam, Compound Sulfonamide, Meropenem, Imipenem, Amikacin, Polymyxin, resistance rates were 98.72%, 98.72%, 96.16%, 96.15%, 96.15%, 66.67%, 60.26%, 58.98%, 58.97%, 57.69%, 57.69%, 57.69%, 57.69%, 57.69%, 56.41%, 55.13%, 53.85%,52.56%, 52.56%, 50%, 0%, see Table 4.

**Tab. 4.** Analysis of drug resistance of *Acinetobacter baumannii*

| Medicine | Drug ResistanceR/% | IntermediaryI/% | Sensitive S/% |
| --- | --- | --- | --- |
| Cefazolin | 98.72 | 0 | 1.28 |
| Ampicillin | 98.72 | 0 | 1.28 |
| Amoxicillin/Clavulanate | 96.16 | 1.28 | 2.56 |
| Aztreonam | 96.15 | 0 | 3.85 |
| Chloramphenicol | 96.15 | 0 | 3.85 |
| Tetracycline | 66.67 | 0 | 33.33 |
| Cefotaxime | 60.26 | 17.95 | 21.79 |
| Ampicillin/Sulbactam | 58.98 | 1.28 | 39.74 |
| Piperacillin | 58.97 | 6.41 | 34.62 |
| Gentamicin | 57.69 | 6.41 | 35.90 |
| Ciprofloxacin | 57.69 | 0 | 42.31 |
| Levofloxacin | 57.69 | 0 | 42.31 |
| Moxifloxacin | 57.69 | 0 | 42.31 |
| Cefepime | 57.69 | 0 | 42.31 |
| Ceftazidime | 56.41 | 2.56 | 41.03 |
| Piperacillin/Tazobactam | 55.13 | 0 | 44.87 |
| Compound Sulfonamide | 53.85 | 0 | 46.15 |
| Meropenem | 52.56 | 0 | 47.44 |
| Imipenem | 52.56 | 0 | 47.44 |
| Amikacin | 50 | 1.28 | 48.72 |
| Polymyxin | 0 | 0 | 100 |

### 2.3 Detection of carbapenem genes in *Acinetobacter baumannii*

Resistance-related genes of *Acinetobacter baumannii bla*_OXA-51_. The electropherograms of PCR products are shown in Figures 1(B). No *bla*_OXA-23_,*bla*_OXA-24_, *bla*_IMP_, *bla*_OXA-58_ and *bla*_SIM_ genes.

**Fig. 1.**
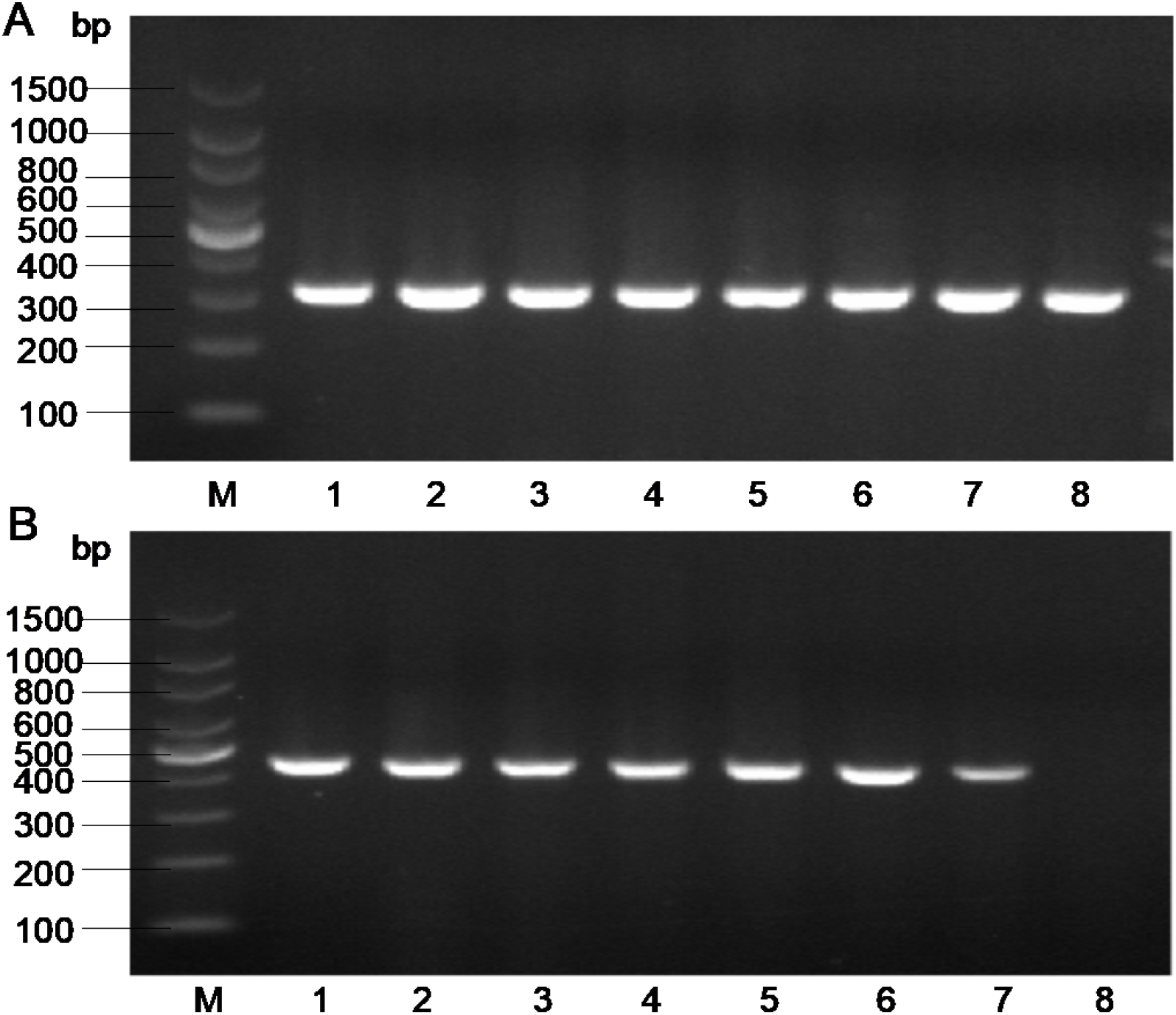
A: Electropherogram of *Acinetobacter baumannii* virulence gene *bfmS* M stands for DNA marker; 1~8 stands for *abaI* geneamplification negative strain;4~6 stands for *bfmS* geneamplification positive strain B: Electropherogram of *Acinetobacter baumannii* drug resistance gene *bla*_OXA-51_ M stands for DNA marker; 1~7 stands for *bla*_OXA-51_ geneamplification positive strain; 8 stands for *bla*_OXA-2_3 geneamplification negative strain

### 2.4 Drug resistance-related gene sequencing

The sequencing results of the PCR products of *Acinetobacter baumannii* drug resistance-related genes *bla*_OXA-51_ are shown in Figures 2(B) and its sequence alignment results shown in Figures 2(A). The comparison in GenBank found that the consistency rates were both above 97%.

**Fig. 2.**
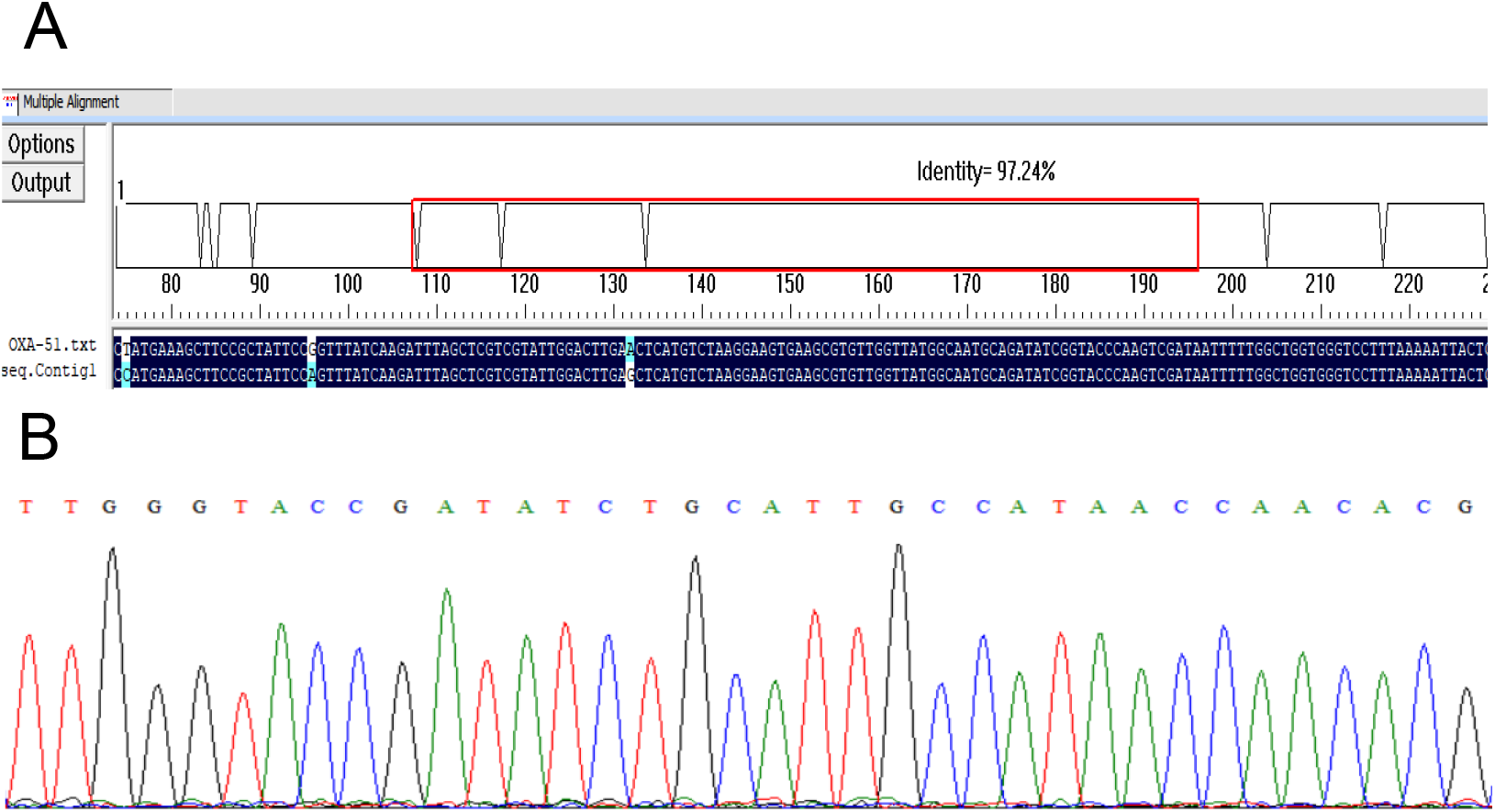
A: *bla*_OXA-51_ gene sequencing alignment B:*bla*_OXA-51_ gene sequencing map

### 2.5 Virulence gene testing

Resistance-related genes of *Acinetobacter baumannii bfmS*. The electropherograms of PCR products are shown in Figures 1(A). No *abaI* and *csuE* genes. The electropherogram of the PCR products of *Acinetobacter baumannii* virulence genes *bfmS* is shown in Figure 3(B) and its sequence alignment results shown in Figures 3(A). The comparison in GenBank found that the consistency rates were both above 97%.

**Fig. 3.**
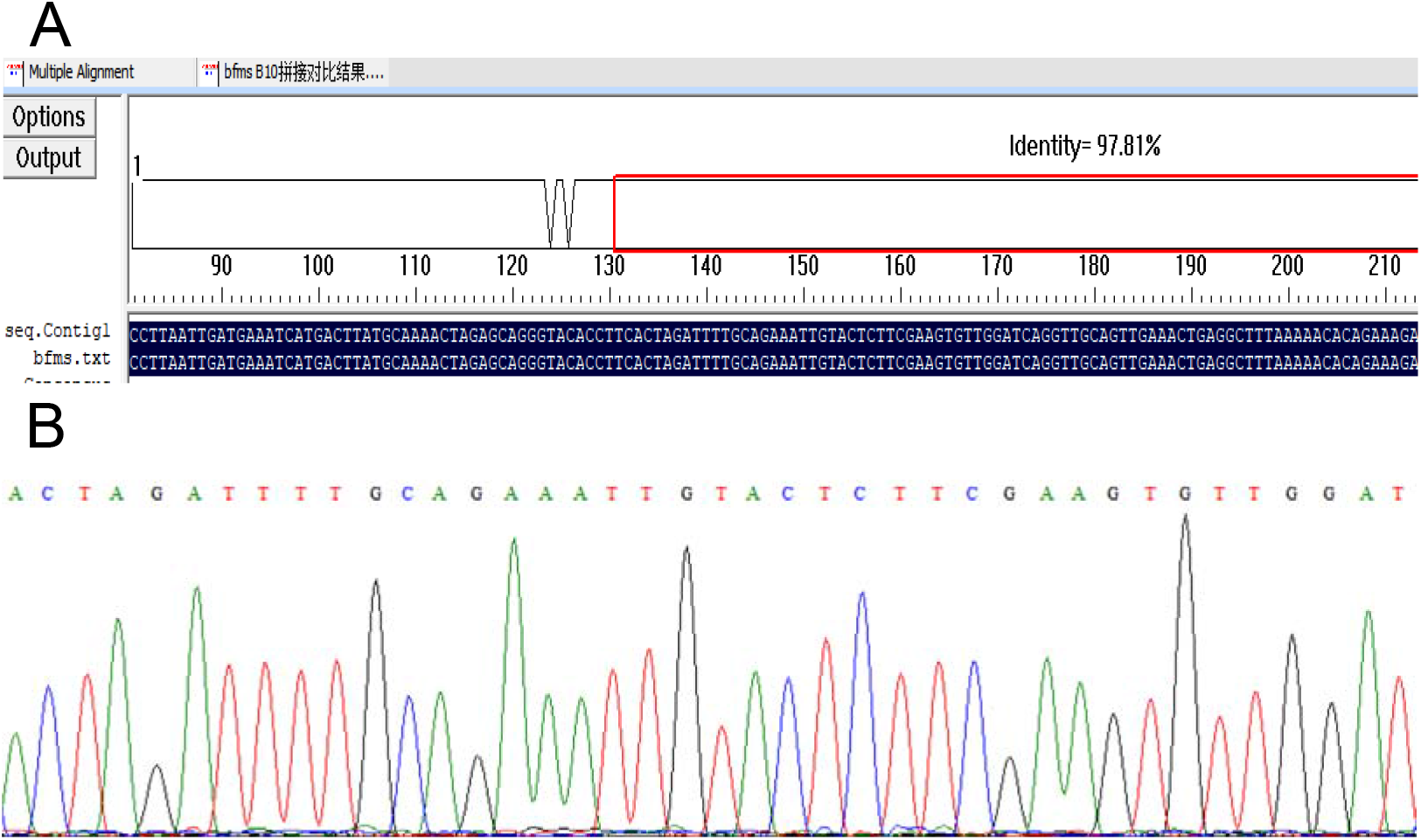
A: Virulence *bfmS* gene sequencing alignment B: Virulence *bfmS* gene sequencing map

## Discussion

In this study, 78 strains of *Acinetobacter baumannii* were sensitive to Cefazolin, Ampicillin, Amoxicillin/Clavulanate, Aztreonam and Chloramphenicol.The resistance rates of Tetracycline and Cefotaxime were all greater than 60%. The drug resistance rates of Ampicillin/Sulbactam, Piperacillin, Gentamicin, Ciprofloxacin, Levofloxacin, Moxifloxacin, Cefepime, Ceftazidime and Piperacillin/Tazobactam were all greater than 55%, while the drug resistance rate to Polymyxin was 0. It can be seen that Polymyxin has good antibacterial activity against *Acinetobacter baumannii*. In clinical anti-infective treatment, the principles of antibacterial treatment of *Acinetobacter baumannii* infection must be followed: comprehensive consideration should be given to the sensitivity of the infectious pathogen, the site and severity of infection, the pathophysiological status of the patient, and the characteristics of antimicrobial drugs. Committed to the research and development of a new generation of antibacterial drugs.

The drug resistance mechanism of clinically isolated carbapenem-resistant *Acinetobacter baumannii* is complex, which is related to the production of carbapenemase, efflux pump genes and integrons gene expression. The drug resistance mechanism is mainly carbapenemase production. Among the carbapenem-resistant *Acinetobacter baumannii* isolates, carbapenem hydrolases are related to metallo-beta-lactamases and *OXA-type* enzymes, although metallo-beta-lactamases have stronger hydrolytic activity, but *OXA-type* enzymes have a more important role in resistance to carbapenems. The *bla*_OXA_ gene is located on the chromosome or plasmid of the bacteria and is usually mediated by integrons or transposons, so it is easier to transfer to other strains. It also has a certain regionality in its distribution range ^[11]^. Studies have pointed out that *bla*_OXA-51_ genes are drug resistance genes frequently detected in carbapenem-resistant *Acinetobacter baumannii*^[12]^. In this study, *bla*_OXA-51_ genes were also detected in carbapenem-resistant *Acinetobacter baumannii* isolates, and their existence was confirmed by sequencing, which should be paid enough attention. Bacterial biofilm is an important cause of bacterial drug resistance. Due to the barrier effect of extracellular polysaccharides, it can reduce the penetration of antimicrobial drugs, thereby resulting in drug resistance ^[13]^. *Acinetobacter baumannii*-related virulence factors play an important role in the formation of biofilms ^[14]^. The sensor kinase *bfmS* regulates the virulence and drug resistance of the cell envelope structure in *Acinetobacter baumannii*^[15]^. Liou et al.^[16]^ showed that insertional inactivation of the *bfmS* gene in *Acinetobacter baumannii* resulted in reduced biofilm formation. The auto-inducible synthase *abaI* is required for biofilm formation and plays an important role in the later stages of biofilm maturation ^[17]^. Studies have shown that the mutation of *abaI* gene can significantly affect the formation of biofilm, and it is more likely to mutate after the overexpression of *abaI* gene to promote the formation of biofilm ^[18]^.Kim et al. ^[19]^ pointed out that the *bfmS* gene regulates drug resistance by controlling the production of outer membrane vesicles in *Acinetobacter baumannii.* This study found that *Acinetobacter baumannii* containing the virulence gene *bfmS* has higher drug resistance, and there is a significant difference compared with the drug resistance without *bfmS* gene. It can be seen that the virulence gene *bfmS* plays an important role in the drug resistance of *Acinetobacter baumannii*. However, the number of strains in this study is relatively small, and the specific conclusions still need to be verified by large sample experiments. In conclusion, *Acinetobacter baumannii* in our hospital is relatively resistant to carbapenems, mainly carrying *bla*_OXA-51_ resistance genes. The drug resistance of *Acinetobacter baumannii* is closely related to the virulence gene *bfmS*. Effective prevention and control measures should be taken to prevent the spread of the drug resistance and virulence genes of *Acinetobacter baumannii* in hospitals.

Bacterial nosocomial infection is now a very important problem in the world, and nosocomial infection caused by non-enteric-negative bacilli such as *Acinetobacter baumannii* is even more embarrassing, so timely medication is of course the best way to reduce nosocomial infection. However, with the extensive use of drugs, bacterial drug resistance is increasing, so this study is devoted to the analysis of drug resistance and virulence genes of carbapenem-resistant *Acinetobacter baumannii* in our hospital, which can provide information for clinical drug use. Some guidelines recommend effective guidance in order to reduce nosocomial infections and bacterial resistance in hospitals. This is also for effective prevention and control of nosocomial infection and nosocomial transmission, and to avoid the continued spread of carbapenem-resistant *Acinetobacter baumannii* in nosocomial areas.

## Acknowledgements

All authors contributed to study conception and design.The authors gratefully thank Prof. Wei for providing the research ideas and Zhenyu Liu for helping complete the experiment.

## Conflict of Interest

All authors disclosed no relevant relationships.

## Funding Source

Funding to pay Open Access publication charges for this article was provided by Chuandong Wei, Department of Laboratory Medicine, Guangxi Health Commission Key Laboratory of Glucose and Lipid Metabolism Disorders, The Second Affiliated Hospital of Guilin Medical University, Guilin.

## Data Availability

The datasets generated during and/or analysed during the current study are available from the corresponding author on reasonable request.

## Highlight

Acinetobacter baumannii has become the main source of nosocomial infections. The overuse of antibiotics has caused Acinetobacter baumannii to develop drug resistance and become “multidrug-resistant Acinetobacter baumannii”

